# Unveiling the neural correlates of habit in the dorsal striatum

**DOI:** 10.1101/2021.04.03.438314

**Authors:** Y. Vandaele, P.H. Janak

**Author notes:** Corresponding author: Youna Vandaele.

## Abstract

We have recently reported sustained inhibition in the dorsomedial striatum (DMS) and sustained excitation in the dorsolateral striatum (DLS) during execution of a lever press sequence in a discrete-trials task promoting habit. This sustained dorsostriatal activity was present early on, and did not clearly change in step with improved performance over ten training sessions. Early onset of sequence-related neural activity could have resulted from rapid habitual learning promoted by presentation of lever cues, predicting reward availability and delivery. To test this hypothesis, we compared DLS and DMS spiking activity in the discrete trials habit-promoting task with two task variants that promote goal-directed behavior. Comparison of the three tasks revealed that mean neuronal spiking activity was generally sustained across the lever press sequence in the task promoting habit and characterized by overall excitation in DLS and inhibition in DMS relative to baseline. In contrast, mean activity differences in DLS and DMS were much less prominent, and most changes occurred transiently around individual lever presses, in the tasks promoting goal-directed behavior. These results indicate that sequence delineation cues, such as the lever cues in these studies, promote habitual behavior and that this habitual behavior is encoded in the striatum by cue-triggered sustained DLS excitation and DMS inhibition that likely reflects cue-elicited behavioral chunking.

## Introduction

Habits are rigid behaviors that over time no longer require planning or deliberation (Dickinson et al., 1995). In contrast to goal-directed behaviors that rely on response-outcome associations, habitual actions are performed in absence of outcome representation, and are therefore insensitive to outcome devaluation or degradation of the instrumental contingency (Balleine et Dickinson, 1998; Dickinson, 1985). By definition, habits are also characterized by higher performance efficiency. Under stable conditions, it has been suggested that actions become “chunked” together into a behavioral unit, facilitating more efficient and consistent execution (Dezfouli et al., 2014; Dezfouli et Balleine, 2013; Graybiel, 1998; Jin et Costa, 2015; Jog et al., 1999; Smith et Graybiel, 2016). Although habit and behavioral chunking are conceptually related, whether they constitute differing expression of a single cognitive process or represent separate cognitive mechanisms remains unclear (Garr et Delamater, 2019).

The dorsal striatum is linked to habit, skill learning, and behavioral chunking with distinct regional specificities (Graybiel, 1998; Graybiel et Grafton, 2015). Functional manipulations show that the dorsomedial striatum (DMS) is involved in goal-directed control early in training whereas the dorsolateral striatum (DLS) is engaged in the formation of habit and skill learning over more extended training (Corbit et al., 2012, 2014; Corbit et Janak, 2016, 2010; Yin et al., 2006, 2005, 2004; Yin et Knowlton, 2006). In rodent neural recording studies, sequence-related activity has been shown to develop in the dorsal striatum as performance improves in operant tasks (Jin et al., 2014; Jin et Costa, 2010; Martiros et al., 2018) and T-maze tasks (Barnes et al., 2005; Smith et Graybiel, 2013; Thorn et al., 2010), with regional differences (Barnes et al., 2005; Thorn et al., 2010). In particular, the development of a task-bracketing activity in the DLS but not the DMS is proposed to reflect behavioral chunking, a process that often coincides with the formation of habit and the disappearance of deliberation (Redish, 2016; Regier et al., 2015; Smith et Graybiel, 2013). We have recently reported striatal regional differences in sequence-related activity during execution of lever press sequences in a task promoting habitual behavior and high performance efficiency (Vandaele et al., 2019, 2017). In this task (termed discrete-trials fixed-ratio 5; DT5), each trial begins by lever insertion, and lever retraction occurs after five consecutive presses, followed by reward delivery. Notably, the activity patterns in DLS and DMS differed from previous studies and did not change during training despite continued improvements in performance (Vandaele et al., 2019). We hypothesize that rapid habitual learning and behavioral chunking is promoted early in training in this DT5 task by the lever insertion and retraction, potentially acting as reward-predictive cues, and is correlated with the early presence of sequence-related activity, observed primarily as sustained inhibitions (DMS) or excitations (DLS), in the dorsal striatum.

To test this hypothesis, we compared DMS and DLS neural activity in the DT5 task with neural activity recorded within two other tasks with fixed-ratio 5 schedules of reinforcement, but in absence of sequence initiation and cessation cues: 1) a standard fixed-ratio 5 (FR5) task, in which each sequence of five lever presses is followed by reward delivery; and, 2) a version of the FR5 task in which mid-sequence reward port checks are penalized, termed the fixed-sequence 5 (FS5) task. In the FR5 task, in contrast to the DT5 task, rats remain sensitive to outcome devaluation, an indication of goal-directed control, and frequently check the reward port before completion of the ratio, resulting in low performance efficiency (Vandaele et al., 2017). Since performance metrics in the FR5 task strongly differ from those of the DT5 task, we also trained rats in the FS5 task mentioned above, in which premature visits of the reward port before completion of the lever press sequence were penalized. Behavior in this task more closely resembles that in the DT5 task and provided an additional comparison group.

Here, we report the replication of previous results showing that lever pressing is habitual in the DT5 task and goal-directed in the FR5 task (Vandaele et al., 2017). Furthermore, we demonstrate that training rats in the FS5 task favors high performance efficiency that remains under goal-directed control. We observed that sequence-related activity in dorsal striatum in the DT5 task is characterized by sustained DLS excitation and DMS inhibition throughout the lever press sequence, as we previously reported (Vandaele et al., 2019). This pattern was specific to the DT5 task and was bracketed by neural responses to the lever insertion and retraction. In contrast, in the FR5 and FS5 tasks, mean dorsostriatal activity during goal-directed behavior was characterized by flat or transient modulations around individual lever presses with no obvious initiation signals, and relatively less excitation in the DLS relative to the DMS. This was especially notable when comparing the FS5 task to the DT5 task, two procedures with relatively similar behavioral profiles. Thus, relatively greater excitation in DLS and sustained inhibition in DMS is correlated with behavioral demonstration of habit. Together these results suggest that lever cues in the DT5 task promote rapid habit learning, encoded in the striatum by sustained DLS excitation and sustained DMS inhibition, whereas the presence of outcome expectations in the FR5 and FS5 tasks results in the absence of this clear population distinction in activity during performance of the five lever press response.

## Results

### Differences in expression of habit and performance efficiency in the DT5, FR5 and FS5 tasks

Rats were trained in three different lever pressing tasks under fixed-ratio 5 schedules of reinforcement (Fig 1A-C) and recordings were obtained from the DMS and DLS (Fig. 1 D,H,L). The first group was trained in the discrete trial fixed-ratio 5 task (DT5 group N=6; Fig. 1A; D-F). In this task, every trial began with insertion of a lever after which rats had to complete a sequence of 5 lever presses to obtain access to a reward, signaled by lever retraction occurring at the fifth lever press (Fig. 1A). Rats quickly learned to complete every ratio of the session as shown by the number of lever presses rapidly reaching the maximum allowed per session (i.e. 150 lever presses; Fig 1E). After 10 DT5 sessions, we confirmed that, on average, responding was insensitive to outcome devaluation induced by sensory-specific satiety (Fig. 1F-G; F_(1,5)_=2.39, p>0.1; devaluation ratio t-test against 0.5: t=1.51, p>0.1).

**Figure 1.**
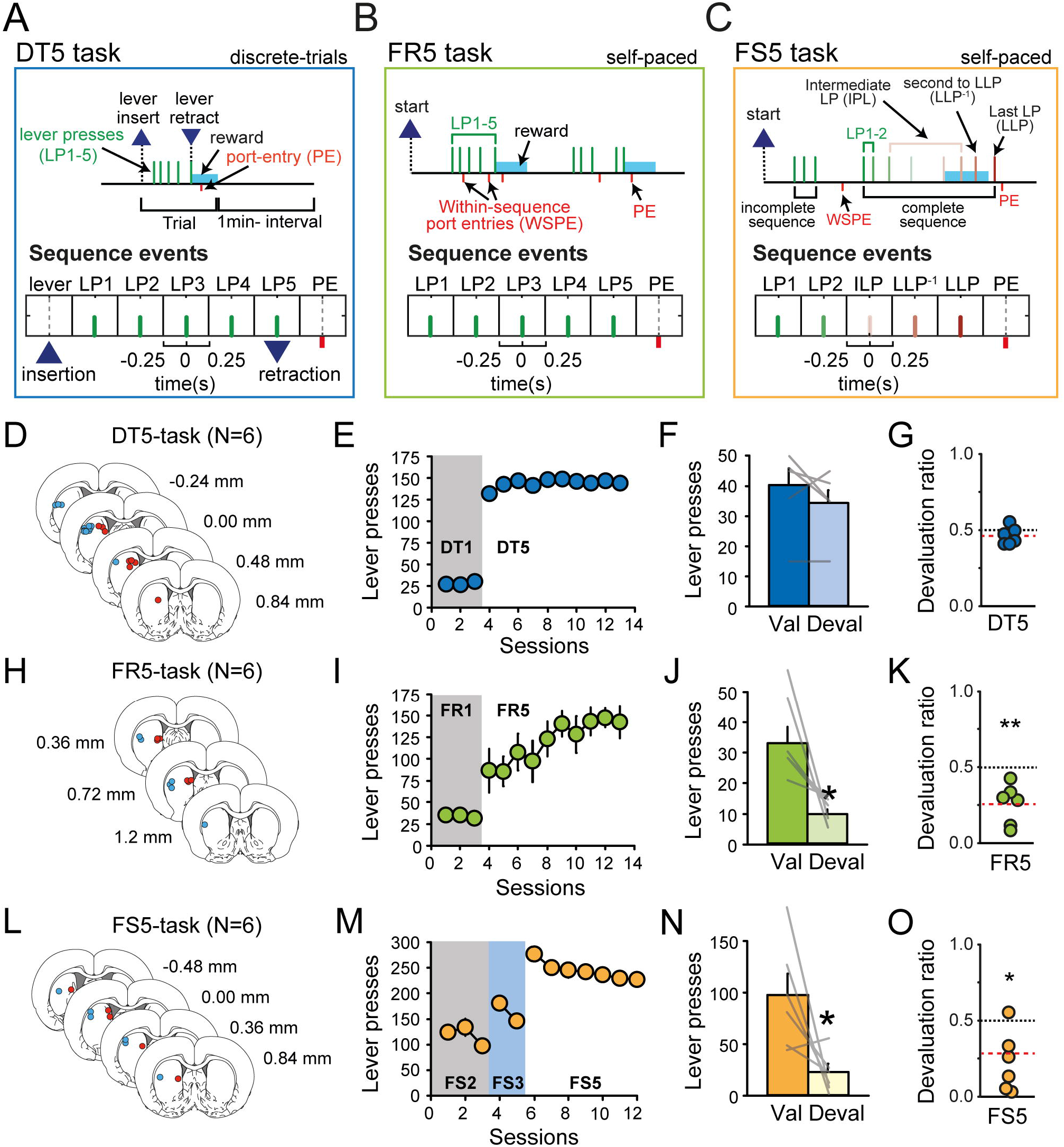
Differences in habitual learning in the DT5, FR5 and FS5 tasks. **A-C**. Diagrams of task - structure and events in the DT5 (A), FR5 (B) and FS5 procedures (C). **D**. Electrode placements in the DT5 group. **E**. Mean number of lever presses across training sessions in the DT5 task. **F**. Mean number of lever presses in the valued and devalued conditions. Grey lines represent individual rats. **G**. Devaluation ratio of individual rats in the DT5 group. Devaluation ratios at or above the black dotted line indicate habit. The red dotted line indicates the group average. **H-K**. Same as D-G for the FR5 group. **L-O**. Same as D-G for the FS5 group. Error bars indicate sem. * p<0.05; ** p<0.01.

The second group was trained in the FR5 task (N=6; Fig. 1B; H-I). In this free-operant task, rats completed 5 lever presses on a lever that was continuously available, and in absence of any reward predictive cue (Fig 1B). The total number of lever presses progressively increased to reach the maximum possible around the last three sessions (Fig. 1I). We confirmed at the end of this training that behavior was goal-directed as indicated by rats’ sensitivity to satiety-induced devaluation (Fig. 1J-K; F_(1,5)_=11.14, p<0.05; devaluation ratio t-test against 0.5: t=4.54; p<0.01).

The third group of rats was trained in the fixed sequence length 5 task (FS5, N=6; Fig 1C; M-O). In this free-operant task, rats had to complete a sequence of at least 5 consecutive lever presses without checking the reward port to obtain a reward (Fig. 1C). A port entry before completion of the ratio reset the ratio and that lever press sequence was considered as incomplete. However, additional presses after completion of the ratio were without consequence and considered as part of the completed sequence (Fig 1C). Importantly, no external cues indicated correct completion of the sequence. After initial training with FS2 and FS3 requirements, rats were tested in the final FS5 schedule for 7 sessions (Fig. 1M; methods). Sensitivity to satiety-induced devaluation was assessed at the end of FS5 training and revealed that behavior was under goal-directed control (Fig. 1N-O; F_(1,5)_=7.48, p<0.05; devaluation ratio t-test against 0.5: t=3.37, p<0.05). Collectively, these results extend previous findings showing that lever pressing is habitual in presence of lever cues signaling the opportunity to respond (DT5 task) but is under goal-directed control in their absence (FR5 and FS5 tasks).

Behavior across the three tasks also differed with respect to sequence learning and performance efficiency. Rats rapidly learned to complete the required ratio in a sequence of consecutive lever presses in the DT5 task by pressing the lever until its retraction on every trial (Fig. 2A). This resulted in suppression of within-sequence port entries by the 2nd session (Fig. 2B; session Friedman ANOVA χ^2^=41.76, p<0.0001), and, across the training period, an increase in within-sequence response rate (Fig. 2B; F_(9,45)_=9.37, p<0.0001) accompanied by a decrease in trial-by-trial variability (Fig. 2B; F_(9,45)_=2.60, p<0.05). In contrast, in absence of lever cues in the FR5-task, rats regularly checked the port after a few lever presses and before completing the ratio (Fig. 2C). This resulted in a high number of within-sequence port entries, a low within-sequence response rate and higher trial-by-trial variability (Fig. 2D; last three sessions DT5 vs FR5: within-sequence port entries Z=2.88, p<0.01; within-sequence response rate F_(1,10)_=45.36, p<0.0001; coefficient of variation Z=2.72, p<0.01). Although within-sequence response rate increased across sessions (session F_(9,36)_=7.97, p<0.0001), it never reached the level observed in the DT5-task (Fig 2D and G). These results suggest that lever insertion and retraction in the DT5 task constitute salient cues that shape behavior, and promote the expression of highly efficient and habitual responding.

**Figure 2:**
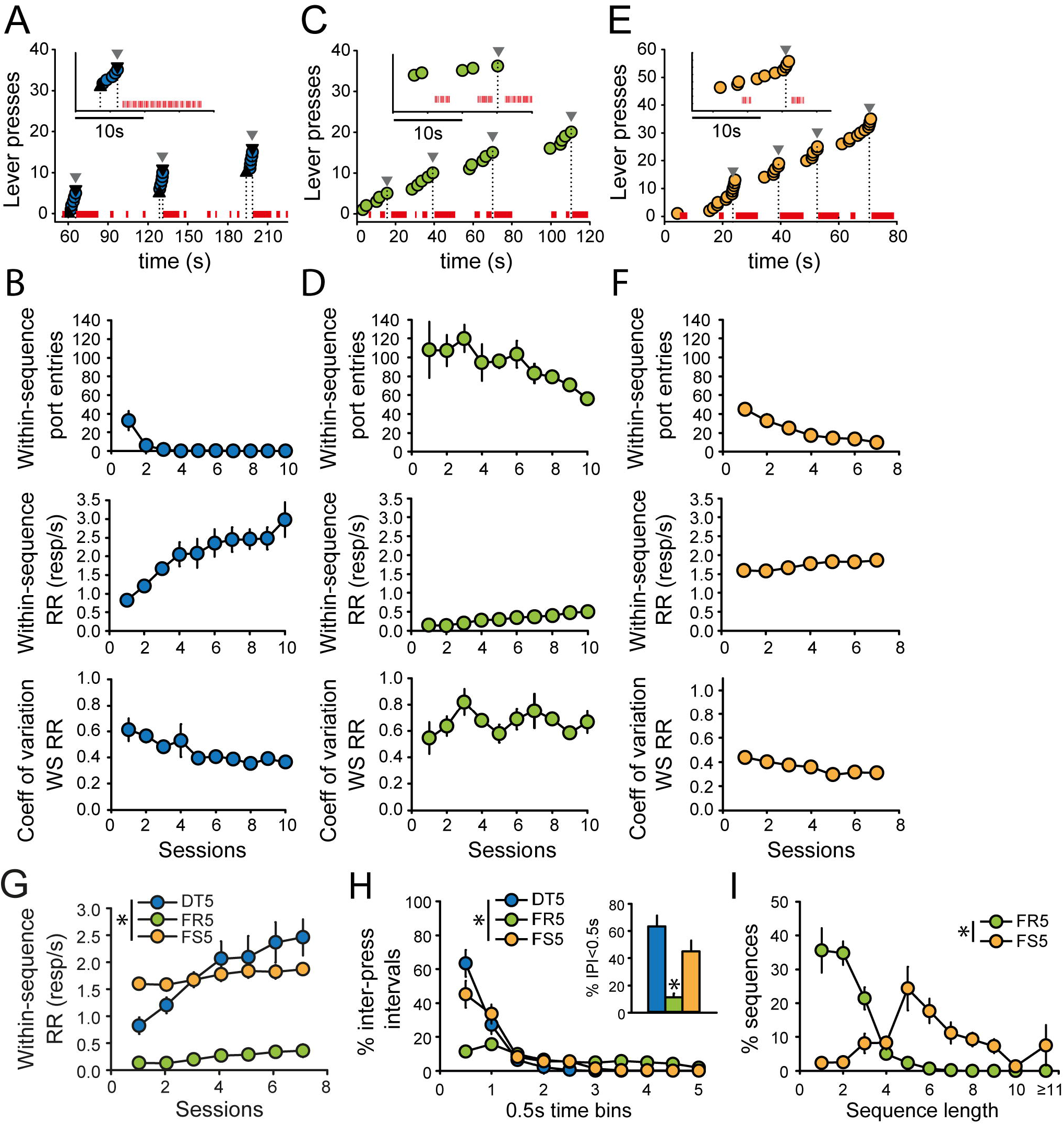
Task differences in performance efficiency. **A**. Microstructure of instrumental performance in a representative rat during DT5 training. Circles represent lever presses; red ticks indicate time in the reward port; upward and downward triangles represent the lever insertion and retraction, respectively. Inset: enlargement of the second trial. **B**. Mean number of within-sequence port entries (top), within-sequence response rate (WS RR; middle) and coefficient of variation of within-sequence response rate (bottom) across DT5 sessions. **C-D**. Same as A-B for the FR5 task. **E-F**. Same as A-B for the FS5 task. In C and E, downward triangles indicate reward delivery. **G**. Mean within-sequence response rate across the first 7 training sessions in the DT5, FR5 and FS5 tasks. *P<0.0001. H. Mean distribution of inter-press intervals (IPI) during the last training session in the DT5, FR5 and FS5 tasks. *P<0.0001. Inset: percentage of IPI shorter than 0.5s in the three tasks. *P<0.001. I. Mean distribution of sequence lengths during the last training session in the FR5 and FS5 tasks. *P<0.0001. Error bars represent the sem.

In line with our predictions, the sequence requirement of the FS5 task greatly improved performance as rats learned to suppress port checking during sequence execution (Fig. 2E-F). The number of within-sequence port entries declined over training and was significantly lower than in the FR5 task (Fig. 2F; main effect of session F_(6,30)_=12.12, p<0.0001; last three sessions FR5 vs FS5: Z=2.88; p<0.01). During complete sequences, the within-sequence response rate was higher than in the FR5 task (main effect of task F_(1,9)_=43.10, p<0.0001) but did not significantly differ from the DT5 task (Fig. 2F-G; FS5 vs DT5: main effect of task F_(1,10)_=0.26, p>0.5) and was associated with similar trial-by-trial variability as the DT5 task (Fig 2F; FS5 vs DT5: main effect of task F_(1,10)_=2.87, p>0.1). Analysis of the distributions of inter-press intervals (IPI) during the last training session in each task revealed that while FR5-performance is characterized by a wide range of IPI, performance in the DT5 and FS5 tasks was characterized by a majority of IPI shorter than 0.5s (Fig. 2H; bins by task interaction F_(18,135)_=10.97, p<0.0001; %short IPI: main effect of task F_(2,15)_=15.95, p<0.001; post-hoc FR5 vs DT5: p<0.001; FR5 vs FS5: p<0.01; DT5 vs FS5: p=0.16). Overall, the performance in the FS5 task was more comparable to the DT5-task but strongly differed from the FR5 task. Although the FR5 and FS5 tasks shared the same free-operant structure with higher uncertainty about reward delivery, rats trained in these 2 tasks adopted very different strategies. More specifically, rats made a majority of short sequences in the FR5-task (1-3 lever presses) whereas performance in the FS5 task was characterized by a wider range of sequence lengths, with a peak at 5 lever presses (Fig. 2I; task*length F_(10,90)_=21.38, p<0.0001). In fact, rats trained in the FS5-task learned to avoid making incomplete sequences by pressing more persistently on the lever, which resulted in a high proportion of sequences longer than 5 lever presses (Fig. 2I; 54.3±10.7%).

### Distinct sequence-related activity in the tasks promoting habitual vs goal-directed behavior

Training in the 3 tasks favored distinct behavioral phenotypes, differing in terms of sensitivity to devaluation and in performance efficiency. We next compared DMS and DLS activity in these 3 tasks to determine how specific activity patterns might relate to instrumental performance and strategy. Medium spiny neurons (MSN) were distinguished from interneuron populations using firing rates and waveform properties (Figure 3-supplement 1A-E) (Martiros et al., 2018; Schmitzer-Torbert et Redish, 2008; Stalnaker et al., 2016). Putative MSNs represented 94%, 87% and 91% of the recorded units in the DT5, FR5 and FS5 groups, respectively. We restricted analyses in this study to the population of putative MSN. Electrode placements were similar across groups (Fig 3-supplement 1). As previously reported (Vandaele et al., 2019), in the DT5-task, task-responsive neurons (TRN) represented on average 86.3±1.8% of the MSN population (DLS, range N=16-34/session; DMS, range N=50-64/session). Average normalized activity of TRN differed as a function of brain region and across events (Fig 3A-D; main effect of region, F_(1,749)_=68.33, p<0.0001; region*event interaction, F_(11,8239)_=25.3, p<0.0001). Specifically, relative to baseline, TRN in DLS expressed sustained excitation throughout the behavioral sequence, from the insertion of the lever to the port entry following reward delivery (Fig 3A). In contrast, DMS activity was characterized by sustained inhibition during lever pressing and excitation at the boundaries of the sequence, at the time of lever insertion and retraction (Fig 3B). Because we did not observe any effect of training session (Fig 3C; F_(9,749)_=0.12, p>0.5), sessions were combined for illustration purpose (Fig 3D) but were systematically considered as a between-factor variable for statistical tests. In the FR5 task, the mean activity of TRNs in DLS and DMS, representing on average 86.9±2.8% of recorded MSNs (DLS, range N=21-43/session; DMS, range N=41-67/session), strongly differed from the DT5 group, with transient excitations around each individual lever press (Fig 3E-H). There was no main effect of region (F_(1,757)_=2.31, p>0.1). DLS and DMS activity significantly differed across events (Fig 3E-F; region*event interaction, F_(11,8327)_=10.64, p<0.0001) but not across sessions (Fig G; main effect of session: F_(9,757)_=0.93, p>0.4; region*session interaction, F_(9,757)_=1.72, p=0.08). While DLS activity peaked at the time of lever presses (Fig. 3E & 3H), DMS activity peaked after each press with higher excitation following the last lever press in the sequence (Fig 3F & 3H). In the FS5-task, the average proportion of TRN across sessions was 90.8±6.1% (DLS, range N=7-27/session; DMS, range N=42-60/session). Again, there was no significant main effect of region (F_(1,466)_=0.13,p>0.5), but an interaction of region x events in this task (Fig. 3I-J; region*event interaction F_(11,5126)_=12.02, p<0.0001). Specifically, mean TRN activity in the DLS transiently decreased after each lever press and increased around the port entry whereas TRN activity in the DMS slowly ramped up toward the end of the lever press sequence (Fig. 3I-L). We did not observe any change in activity across training sessions (Fig. 3K; F_(6,466)_=0.93, p>0.4).

**Figure 3.**
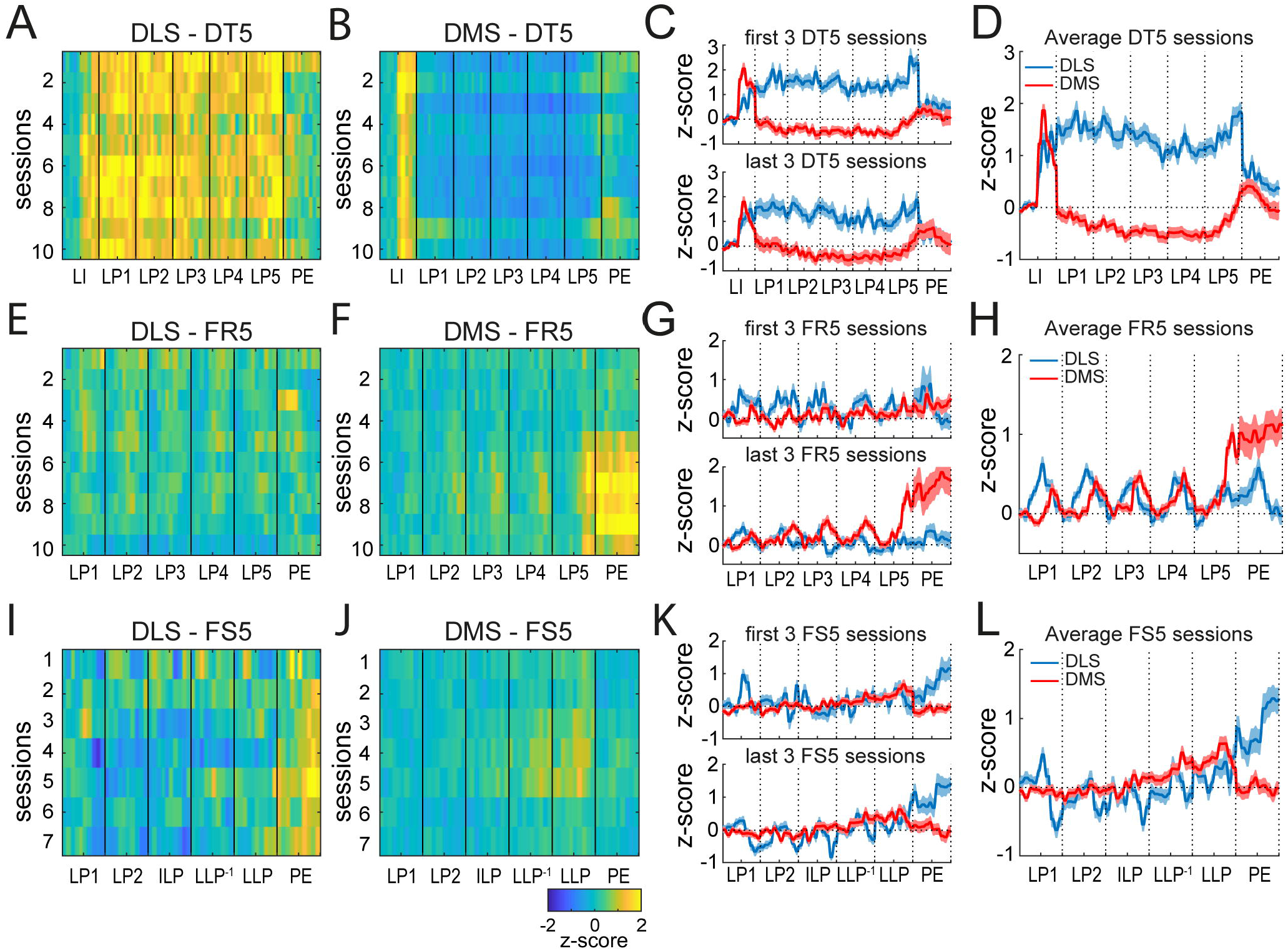
Distinctions among sequence-related activity in the DT5, FR5 and FS5 tasks. **A-B**. Heatmaps of average TRN activity along the behavioral sequence and across training sessions in the DLS (A) and DMS (B). **C**. Average PSTH of DLS and DMS TRN over the first three (top) and last three (bottom) DT5 training sessions. **D**. Average PSTH of DLS and DMS TRN combined across all training sessions. **E-H**. Same as A-D for the FR5 task. **I-L**. Same as A-D for the FS5 task.

The above analysis demonstrates distinct patterns of activity as a function of region in each task with a distinct lack of an overall mean difference in firing between DLS and DMS in the two tasks with goal-directed behavior. We next asked if neural activity patterns differed among the three tasks using a statistical comparison. A 3-way ANOVA with task, region and session as between-factors (analysis restricted to first 7 sessions) and task event as a repeated factor, revealed significant effects of task (F_(2,1425)_=21.55, p<0.0001), and a task by region (F_(2,1492)_=6.14, p<0.01) and task by region by event (F_(22,16412)_=11.25, p<0.0001) interactions. Post hoc analysis revealed significant differences in DLS and DMS activity between the DT5 and the other two tasks (p-values<0.0001) but not between the FR5 and FS5 tasks (p-values>0.3). Together, these results reveal differences in mean activity in the DLS and DMS that distinguish among the three tasks, with a relatively greater inhibition in DMS present in the DT5 task in which behavior is habitual.

### Persistent task differences in activity pattern after controlling for difference in performance efficiency

The above analyses revealed important task differences in activity pattern that could be attributed to differences in habit and behavioral chunking or performance efficiency. Indeed, although the DT5 task is the only task to promote habit and lever-triggered behavioral chunking, it is also the task associated with the highest performance. Although instrumental performance in the FS5 task was comparable to the DT5 task, rats remained more efficient in the DT5 task and tended to execute lever press sequences with very high response rates. To control for differences in response rate between the DT5 and FS5 tasks, for each recording session, we matched trials from the DT5 dataset to the middle tertile of response rates in the FS5 dataset (Fig 4A). Using this approach, within-sequence response rate was overall higher in the FS5 task than in subsampled trials in the DT5 task (Fig 4B; main effect of task: F_(1,1332)_=5.56, p<0.05). However, session by session analysis revealed no task difference (F-values<2.19, p-values>0.1).

**Figure 4:**
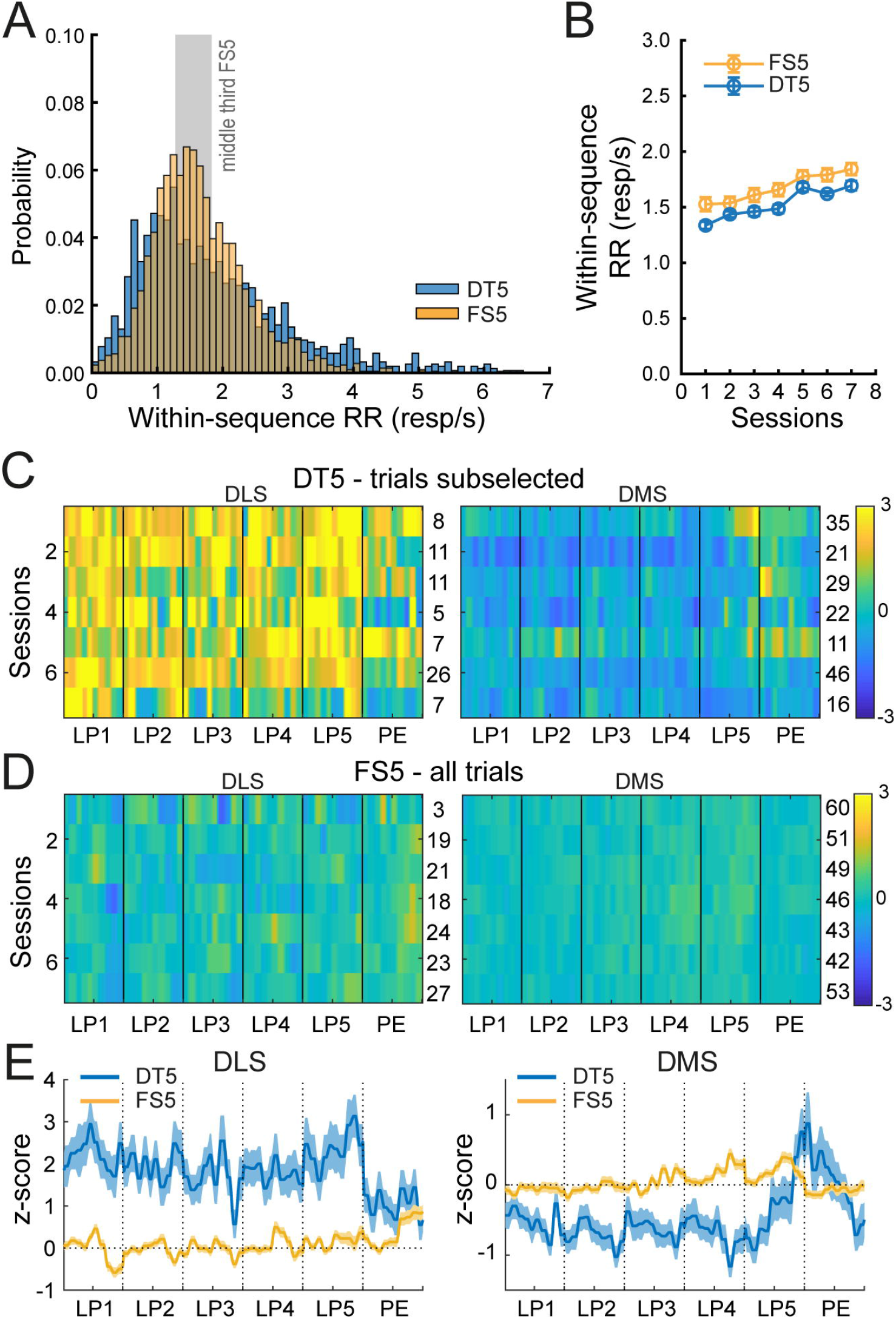
Task differences in activity pattern despite similar within-sequence response rate. **A**. Distribution of within-sequence response rates across recording sessions in the DT5 and FS5 tasks. The grey area represents the middle tertile of within-sequence response rates in the FS5 task, in which we selected trials from the DT5 task. **B**. Mean within-sequence response rate (±sem) across training sessions in the DT5 and FS5 tasks after subsampling of DT5 trials. **C-D**. Heatmaps of the average TRN activity in DLS (left) and DMS (right) across sessions in the DT5 task after trials subsampling (C) and in the FS5 task (D). Numbers on the right of the heatmaps indicate the number of neurons. **E**. Average PSTH (±sem) of DLS (left) and DMS (right) TRN in the FS5 task and in the DT5 task after trial subsampling.

After controlling for fast sequences by restricting our analysis of DT5 activity to the middle range of FS5 response rates, we observed that the average TRN activity in DLS and DMS remained significantly different from the FS5 task (Fig 4C-F). A 3-way ANOVA with task, region and session as between-factors and event as a repeated factor revealed a main effect of task (F_(1,691)_=42.81, p<0.0001), a task by region interaction (F_(1,691)_=132.18, p<0.0001), a task by event interaction (F_(11,7601)_=4.72, p<0.001) and a task by region by event interaction (F_(11,7601)_=11.20, p<0.0001). Compared to the FS5 task, the average TRN activity in the DT5 task showed a sustained excitation in the DLS (Fig 4C and E) and a sustained inhibition in the DMS (Fig 4D and F), while the mean activity in both the DLS and DMS during the FS5 task tended to return to baseline after any deviation during the five lever-press responses. Thus, neuronal activity patterns in the DT5 task significantly differed from the FS5 task despite similar or lower response rates. These results suggest that higher performance efficiency alone cannot explain the sequence-related activity pattern of sustained inhibition or excitation observed in the DT5 task.

## Discussion

In this study, we report distinctions in striatal neural activity in 3 different tasks sharing the same response requirement but differing in terms of instrumental performance and strategy. The DT5 task was the only task promoting habitual responding with high performance efficiency by providing discrete lever cues predicting reward availability and delivery. In the DT5 task, we previously observed rapid development of distinct sequence-related activity patterns in the DLS and DMS, characterized by sustained excitation throughout the sequence in the DLS and sustained inhibition during lever presses with excitations at the sequence boundaries in the DMS. In clear contrast, dorsostriatal activity during goal-directed behavior in the FR5 and FS5 tasks was characterized by a lack of an overall mean difference in DMS and DLS activity, and by more transient modulations around individual lever presses that typically returned to baseline. Together these results suggest that lever-triggered sequence-related activity in dorsal striatum emerging from the very beginning of training in the DT5 task directly encodes habit, promoted by the lever insertion and retraction cues provided in the task.

We replicated previous results showing that the presence of lever cues in the DT5 task promoted habitual learning with high performance efficiency whereas their absence in the FR5 task resulted in a goal-directed behavior with low performance efficiency (Vandaele et al., 2017). We are now extending these results, by showing that training in the FS5 task allowed for sequence learning but did not promote habitual learning. In the FR5 and FS5 tasks, the lever was continuously extended and could not constitute a cue predictive of reward availability or delivery. The high number of within-sequence port entries in the FR5 task and the rats’ tendency to keep pressing on the lever after the reward delivery in the FS5 task reveals the uncertainty associated with the reward delivery in these conditions. It was recently suggested that high reinforcer predictability could favor the development of habit by reducing the attentional demand on behavioral monitoring (Thrailkill et al., 2018; Vandaele et al., 2020; Vandaele et Ahmed, 2020). This could explain the rapid development of habit, and perhaps behavioral chunking, in the DT5 task, in which the lever retraction directly signals the completion of the sequence and can ameliorate any requirement for monitoring the number of lever press responses. In contrast, the sequence requirement and the uncertainty about reward delivery in the FS5 task may require rats to maintain a representation of the outcome by keeping track of the time elapsed or the number of lever presses made before checking the port. This cognitive demand may prevent the development of habit and explain rats’ sensitivity to outcome devaluation (Kosaki et Dickinson, 2010). As previously suggested (Garr et Delamater, 2019), these results show that performing an action sequence with great automaticity does not imply that responding is habitual and, by extension, that goal-directed control should not be equated with a lack of automaticity.

The role of dorsal striatum in motor control and skill learning raise the possibility that task differences in dorsostriatal activity reflect differences in performance efficiency rather than habitual behavior *per se* (Dudman et Krakauer, 2016; Kupferschmidt et al., 2017; Robbe, 2018; Rueda-Orozco et Robbe, 2015; Sales-carbonell et al., 2018). Although instrumental performance in the FR5 task strongly differed from the DT5 task, the analysis of complete sequences in the FS5 task indicates a performance more comparable with similar within-sequence response rate during several sessions (i.e. sessions 3-5) and low trial-by-trial variability in this measure (Fig 2). Furthermore, the activity patterns in DLS and DMS significantly differed between the DT5 and FS5 tasks after controlling for session-by-session differences in response rate (Fig 5), suggesting that the difference in performance efficiency is not sufficient to explain the results reported here. Importantly, differences in activity pattern cannot be attributed to differences in electrode placements across task, or to differences in response requirement, reward exposure or training duration. However, different reward contingencies may have been experienced in the FS5 task, which, by its nature, resulted in a larger proportion of unrewarded lever presses due to incomplete sequences and extra-lever presses in complete sequences. Further research is needed to determine whether and to what extent the perceived reward contingency may have affected striatal activity.

The average activity of TRN in DMS and DLS strongly differed across task. In the DT5-task, we observed dissociated sequence-related activity in DMS and DLS characterized by sustained excitation throughout the behavioral sequence in the DLS, and a sustained inhibition during lever pressing with excitations at the boundaries of the sequences in the DMS (Vandaele et al., 2019). In marked contrast, mean modulation of spiking activity in the FR5 and FS5 tasks was much less distinct across regions. While DLS neurons in the FR5 task were mostly modulated around lever presses, DMS neurons peaked after every lever press with a stronger activity at the termination of the sequence, notably during the last 5 sessions. In the FS5 task, neural activity in DMS tended to ramp up as rats progressed in the execution of the sequence. This ramp-like pattern may reflect timing processes or a rise in expectation with the approach of the reward. Interestingly, these results differ from a previous study showing start-stop responses and sustained excitation or inhibition during execution of fast lever press sequences with a time requirement in mice (Jin et al., 2014). Differences in species, behavioral control, instrumental constraint (time versus sequence requirement) and/or in the occurrence of behavioral chunking may explain this discrepancy.

In the FR5 and FS5 tasks, excited TRN in DLS peaked around each individual lever press whereas the peaks of activity in DMS neurons occurred after the lever presses, with larger responses following the last lever press, when reward is delivered. These results suggest that while DLS activity is more closely related to the execution of actions, peaks in DMS activity may represent outcome expectation. In line with this idea, a peak in DMS activity is only observed after the last lever press in the DT5 task, when reward delivery is imminent and predicted by the lever retraction, but occurs after each individual lever presses during the FR5 and FS5 tasks, in which there is uncertainty about the time of reward delivery. According to this hypothesis, the activity pattern of DMS neurons in the DT5 task, characterized by sustained inhibition during lever presses followed by an excitation at the termination of the sequence, could be understood as a neural correlate of action chunking; that is, individual actions are executed as a behavioral unit without outcome representation until the completion of the sequence.

Behavioral chunking may also explain why the activity of DLS neurons is sustained across the entire lever press sequence in the DT5 task, but shows more transient modulations around individual presses in the FR5 and FS5 tasks. In the dorsal striatum, the increase in activity generally precede the initiation of movement and peak during the execution of actions (Apicella et al., 1992; Hassani et al., 2001; Hikosaka et al., 1989; Hollerman et al., 1998; Kimchi et al., 2009; Pasupathy et Miller, 2005). However, specific patterns of activity including start/stop or sustained activity may emerge when multiple actions get concatenated to form a single sequence (Jin et al., 2014; Jin et Costa, 2015; Smith et Graybiel, 2016), like in the DT5 task (Vandaele et al., 2019). Although we cannot exclude that sustained DLS excitation in the DT5 task results from shorter inter-press intervals, dissimilar activity patterns in the FS5 and DT5 tasks when differences in response rate are controlled for, suggest otherwise. Thus, it is tempting to speculate from the comparison of dorsostriatal activity across the three tasks that although rats are constrained to execute sequences of consecutive lever presses in the FS5 task, they only learn to concatenate individual action into unitary sequences in the DT5 task. Rapid behavioral chunking in the DT5 task (occurring within the 1^st^ session) and its presumed absence in the FR5 and FS5 tasks, could explain why striatal activity do not appear to track improved task performance across training sessions in our conditions.

## Materials and methods

### Subjects

#### Subjects

18 male Long Evan rats (Envigo) were individually housed in a temperature (21° C) and light-controlled (12-h light-dark cycle, lights ON at 7am) vivarium, with partial enrichment. All experiments were performed during the light cycle. Rats were given free access to water throughout the experiment and were maintained at 90% of their free-feeding weight. This study was carried out in accordance with the recommendations of the Guide for the Care and Use of Laboratory Animals (Institute of Instrumental Training Laboratory Animal Resources, Commission of Life Sciences, National Research Council, 1996). The protocol was approved by the institutional animal care and use committee of Johns Hopkins University.

#### Experimental groups

Following instrumental training under continuous reinforcement schedules, rats in the 3 different groups received surgery and neurons in dorsal striatum were recorded throughout acquisition of the DT5 task (N=6), the FR5 task (N=6) or the FS5 task (N=6). Sensitivity to satiety-induced devaluation was assessed at the end of recording in all the rats, tethered in the recording chamber.

### Behavioral training

#### Initial instrumental training

Rats were trained with a small aliquot of a solution of 20% sucrose (0.1mL delivered over 3s). In each task, the house-light, located on the ceiling of the operant chamber remained illuminated during sessions. After a single 30-min magazine training session under a variable interval 60s schedule, rats were trained to press the left lever to earn the reward, delivered in the adjacent magazine. Sessions were limited to 1h or 30 reward deliveries. Rats were trained for 3 to 5 sessions, until they earned the 30 rewards in less than 1h. Conditioning chambers housed within sound-attenuating boxes (Med Associates, St Albans, VT) and designed for in vivo neural recording were used for all training and testing.

#### DT5 task training

Each trial (max: 30) of the session begin with the insertion of the left lever and completion of the lever press requirement results in the retraction of the lever, the reward delivery, and the initiation of a new inter-trial interval. The response requirement was 1 for 3 sessions (discrete-trial FR1; DT1) and increased to 5 for 10 sessions (discrete-trial FR5; DT5). Failure to complete the ratio within 1 minute was considered as an omission and resulted in lever retraction and initiation of a new inter-trial interval. Data from this group of rats were presented in a previous study aimed at comparing DMS and DLS activity across early and extended training in the DT5 task (Vandaele et al., 2019). Here we only selected rats from the early training group trained with liquid 20% sucrose.

### FR5 task training

In this free-operant task, the lever was continuously presented and 5 presses on the lever resulted in a 3-s delivery of the liquid sucrose reward (Fixed ratio 5, FR5). Rats were trained for ten FR5 sessions, limited to 30min or 30 reward deliveries.

### FS5 task training

In this free-operant task, the lever was continuously presented. Rats had to complete a sequence of at least 5 consecutive lever presses without checking the port to obtain a reward, whose delivery was not signaled. A port entry before completion of the ratio resulted in resetting the ratio and the lever press sequence was considered as incomplete. However, additional presses after completion of the ratio were without consequences and considered as part of the completed sequence. Rats were first trained with a fixed sequence length of 2 lever presses for a minimum of 3 sessions or until they earned 30 rewards. The response requirement was then increased to 3 lever presses for a minimum of 2 sessions (or until earning 30 rewards) before training in the final fixed sequence length 5 schedule (FS5) for 7 sessions. Sessions were limited to 30min or 30 reward deliveries.

#### Outcome devaluation by sensory-specific satiety

Each rat received 2 days of testing, separated by one reinforced training session. Rats were given 1h free access to their training reward (sucrose 20%; devalued condition) or to a control reward, which never served as a reinforcer (grain-based pellet; valued condition). Pre-feeding occurred in feeding cages in the experimental room. Immediately after pre-feeding, rats were placed in the recording chambers for a test session conducted under extinction. Test sessions were limited to 10 trials in the DT5 task and 10 minutes in the FR5 and FS5 tasks. On the second test session, animals were pre-fed with the alternative reward prior to the extinction test.

### Electrophysiological recordings

#### Surgeries and recording

After acquisition of instrumental responding, rats underwent surgeries. In the DT5 group, rats were implanted with 2 unilateral arrays of 8 wires aimed at DMS and DLS (0.004’ steel wires arranged in a 2 x 4 configuration, each array spaced 2 mm apart, Microprobes). In the FR5 and FS5 groups, 16 tungsten wires were soldered on two connectors and arranged in 2 bundles spaced 2mm apart. For each group, target coordinates were +0.25 mm AP, ±2.3 mm ML, -4.6 mm DV for DMS and +0.25 mm AP, ±4.3 mm ML, -4.6 mm DV for DLS. Surgeries were performed under isoflurane anesthesia (0.5-5%) with pre-operative injections of cefazolin (75 mg/kg, antibiotic) and carprofen (5 mg/kg, analgesic). Topical lidocaine was applied for local analgesia.

After a minimum of 5 days post-operative recovery, rats were accustomed to tethering with the recording cable for a few CRF sessions before the recording began. Cables were connected at one end to rats’ headsets and at the other end to a commutator allowing free movement throughout acquisition of single-unit activity during the recording sessions. The multichannel acquisition processor (MAP) neural recording system (Plexon Inc, TX) was used to store and process amplified signals and timestamps of behavioral events.

### Analysis of electrophysiological recordings

#### Spike sorting

As previously described (Ottenheimer et al., 2018; Richard et al., 2018), individual units were isolated offline using Offline Sorter (Plexon Inc, TX). Analysis of interspike intervals distribution, auto-correlograms and cross-correlograms (Offline Sorter v3 and Neuro-Explorer 3.0, Plexon Inc, TX) was conducted to control for correct isolation of single units. Average waveforms and timestamps of units and event were exported from Neuro-Explorer 3.0 to Matlab (MathWorks, MA) for further analysis. Analyses were restricted to units with well-defined waveforms and consistent characteristics throughout the entire recording session.

#### Waveform analysis

Neurons were classified as putative Medium Spiny Neurons (MSN), Fast Spiking Interneurons (FSI) or Tonically Active Neurons (TAN), based on the half-valley width of the average waveform and the overall firing rate (Figure 3-supplement 1A-E). Putative-FSI were defined by high firing rate (>20Hz) and narrow waveforms (<0.15ms) whereas putative-TAN were defined by low firing rate (<5Hz) and wide waveform (>0.45ms) as previously reported (Martiros et al., 2018; Schmitzer-Torbert et Redish, 2008; Stalnaker et al., 2016). Neurons not classified as interneurons but showing features intermediate to MSNs and interneurons were unclassified (firing rate between 12.5 and 20Hz and half-width between 0.4 and 0.45ms). All the analyses in this study were conducted on the population of putative-MSNs.

#### Characterization of task-responsive neurons and z-scores

In each task, we analyzed spiking activity across twelve 0.25s periods before and after each lever press and port entry events. Since the performance in the FS5 task was characterized by a large proportion of sequences longer than 5 lever presses, the lever press events considered for the analysis were defined as follow: the 1^st^ and 2^nd^ lever presses, one randomly selected intermediate lever press, the second to last lever press and the last lever press.

For each neuron, the presence of significant excitation or inhibition in response to an event was detected by running a t-test on the firing rate during the 0.25s periods pre- and post-event in comparison to a 1-s baseline period beginning 1-s before trial onset (defined by the lever insertion in the DT5 task and by the 1^st^ lever press in the FR5 and FS5 tasks). Neurons expressing a significant response (p<0.01) to at least one of the 12 behavioral events were considered as task responsive neurons (TRN).

Neural activity for each individual neuron was z-scored as follow: (F_i_-F_mean_)/F_sd_, where F_sd_ and F_mean_ represent the standard deviation and mean of the firing rate during the 1-s baseline period, and F_i_ is the firing rate at the i^th^ bin of the peristimulus time histogram (PSTH). Heatmaps and average PSTH presented in this study represent the z-scores from -0.25s to 0.25s around each event of the behavioral sequence.

In figure 5, to control for differences in response rate between the DT5 and FS5 tasks, we matched trials from the DT5 dataset to the middle tertile of response rates in the FS5 dataset. Specifically, for each FS5 sessions, we determined the 33rd and 66^th^ percentile of within-sequence response rates across subjects and trials. On each DT5 recording session and until the 7^th^ session, we restricted the analysis of neuronal spiking activity to trials with response rates between the 33rd and 66^th^ percentile of the FS5 response rates distribution. We only considered DT5 sessions comprising at least 5 trials in this response rate range.

#### Statistical analysis

Within-sequence response rate (in resp/s) was computed by dividing the number of lever presses per sequence by the time from the first to the last lever press. The devaluation ratio is defined by the ratio of the number of lever presses in the devalued condition divided by the total number of lever presses in the valued and devalued conditions. Thus, a devaluation ratio of 0.5 indicates habitual responding. In the FS5 task, only complete sequences were considered in behavioral and electrophysiological analyses. Data following a normal distribution were subjected to repeated measures analysis of variance. The Mann Whitney test was used when normality assumptions were violated. Mean z-scores of units from DMS and DLS were compared across 12 consecutives events using repeated-measures ANOVA (with Geisser-Greenhouse correction for violation of sphericity), with events as within-design factor and regions, sessions and, when relevant, task, as between-design factors. Significant interactions were analyzed using the HSD Tukey post-hoc test. Events consisted of the average z-score during 0.25s periods before and after each lever press and port entry events. Proportions of neurons were compared using chi-squared tests. All analyses were conducted on MATLAB (MathWorks) and Statistica (StatSoft 7.0).

### Histology

Electrode sites were labeled by passing a DC current through each electrode, under deep anesthesia with pentobarbital. All rats were perfused intracardially with 0.9% saline followed by 4% paraformaldehyde (FR5 and FS5 groups with tungsten wires), or 4% paraformaldehyde with 3% potassium ferricyanide (DT5 group with steel wires). Brains were extracted, post-fixed in 4% paraformaldehyde for 4–24hr, and transferred in 20% sucrose for >48 h for cryo-protection. To verify the electrodes placement, the brains were sectioned at 50 µm on a cryostat and slices were stained with cresyl violet and analyzed using light microscopy.

## Acknowledgements

This work was supported by the National Institute for Health Research (R01DA035943 to P.H.J), the National Institute on Alcohol Abuse and Alcoholism (R01AA026306 to P.H.J), and the Peter and Traudl Engelhorn foundation (to Y.V.)

## Author Contributions

Conceptualization, Y.V. and P.H.J.; Methodology, Y.V., P.H.J.; Formal analysis, Y.V. and P.H.J.; Investigation, Y.V.; Writing – Original Draft, Y.V. and P.H.J.; Writing – Review &Editing, Y.V. and P.H.J.; Visualization, Y.V.; Funding Acquisition, P.H.J.; Resources, P.H.J.; Supervision, P.H.J.

## Declaration of Interests

The authors declare no competing interests.

## Notes

### Competing Interest Statement

The authors have declared no competing interest.

## Bibliography

Apicella P, Scarnati E, Ljungberg T, Schultz W. 1992. Neuronal Activity in Monkey Striatum Related to the Expectation of Predictable Environmental. J Neurophysiol 68:945–960.

Balleine BW, Dickinson A. 1998. Goal-directed instrumental action: Contingency and incentive learning and their cortical substrates. Neuropharmacology 37:407–419. doi:10.1016/S0028-3908(98)00033-1

Barnes TD, Kubota Y, Hu D, Jin DZ, Graybiel AM. 2005. Activity of striatal neurons reflects dynamic encoding and recoding of procedural memories. Nature 437:1158–1161. doi:10.1038/nature04053

Corbit L, Nie H, Janak P. 2012. Habitual alcohol seeking: time course and the contribution of subregions of the dorsal striatum. Biol Psychiatry 72:389–395.

Corbit LH, Janak PH. 2016. Habitual Alcohol Seeking: Neural Bases and Possible Relations to Alcohol Use Disorders. Alcohol Clin Exp Res 40:1380–1389.

Corbit LH, Janak PH. 2010. Posterior dorsomedial striatum is critical for both selective instrumental and Pavlovian reward learning. Eur J Neurosci 31:1312–1321. doi:10.1111/j.1460-9568.2010.07153.x.Posterior

Corbit LH, Nie H, Janak PH. 2014. Habitual responding for alcohol depends upon both AMPA and D2 receptor signaling in the dorsolateral striatum. Front Behav Neurosci 8.

Dezfouli A, Balleine BW. 2013. Actions, action sequences and habits: evidence that goal-directed and habitual action control are hierarchically organized. PLoS Comput Biol 9:e1003364.

Dezfouli A, Lingawi NW, Balleine BW. 2014. Habits as action sequences: hierarchical action control and changes in outcome value. Philos Trans R Soc L B Biol Sci 369.

Dickinson A. 1985. Actions and Habits: The Development of Behavioural Autonomy. Philos Trans R Soc B Biol Sci. doi:10.1098/rstb.1985.0010

Dickinson A, Balleine BW, Watt A, Gonzalez F, Boakes RA. 1995. Motivational control after extended instrumental training. Anim Learn Behav 23:197–206. doi:10.3758/BF03199935

Dudman JT, Krakauer JW. 2016. The basal ganglia: From motor commands to the control of vigor. Curr Opin Neurobiol 37:158–166. doi:10.1016/j.conb.2016.02.005

Garr E, Delamater AR. 2019. Exploring the relationship between actions, habits, and automaticity in an action sequence task. Learn Mem 26:128–132. doi:10.1101/lm.048645.118

Graybiel AM. 1998. The Basal Ganglia and Chunking of Action Repertoires. Neurobiol Learn Mem 136:119–136.

Graybiel AM, Grafton ST. 2015. The Striatum: Where Skills and Habits Meet. Cold Spring Harb Perspect Biol 7:a021691. doi:10.1101/cshperspect.a021691

Hassani Oumk, Cromwell HC, Schultz W. 2001. Influence of Expectation of Different Rewards on Behavior-Related Neuronal Activity in the Striatum. J Neurophysiol 85:2477–2489.

Hikosaka O, Sakamoto M, Usui S. 1989. Functional Properties of Monkey Caudate Neurons III.Activities Related to Expectation of Target and Reward. J Neurophysiol 61:814–832.

Hollerman JR, Tremblay L, Schultz W. 1998. Influence of Reward Expectation on Behavior-Related Neuronal Activity in Primate Striatum. J Neurophysiol 80:947–963.

Jin X, Costa R. 2015. Shaping action sequences in basal ganglia circuits. Curr Opin Neurobiol 33:188–196.

Jin X, Costa RM. 2010. Start/stop signals emerge in nigrostriatal circuits during sequence learning. Nature 466:457–462.

Jin X, Tecuapetla F, Costa RM. 2014. Basal ganglia subcircuits distinctively encode the parsing and concatenation of action sequences. Nat Neurosci 17:423–430.

Jog MS, Kubota Y, Connolly CI, Hillegaart V, Graybiel a M. 1999. Building neural representations of habits. Science 286:1745–1749. doi:10.1126/science.286.5445.1745

Kimchi EY, Torregrossa MM, Taylor JR, Laubach M. 2009. Neuronal Correlates of Instrumental Learning in the Dorsal Striatum. J Neurophysiol 102:475–489. doi:10.1152/jn.00262.2009.

Kosaki Y, Dickinson A. 2010. Choice and contingency in the development of behavioral autonomy during instrumental conditioning. J Exp Psychol Anim Behav Process 36:334–342. doi:10.1037/a0016887

Kupferschmidt DA, Juczewski K, Cui G, Johnson KA, Lovinger DM. 2017. Parallel, but Dissociable, Processing in Discrete Corticostriatal Inputs Encodes Skill Learning. Neuron 96:476-489.e5. doi:10.1016/j.neuron.2017.09.040

Martiros N, Burgess AA, Graybiel AM. 2018. Inversely Active Striatal Projection Neurons and Interneurons Selectively Delimit Useful Behavioral Sequences. Curr Biol 28:560-573.e5. doi:10.1016/j.cub.2018.01.031

Ottenheimer D, Richard JM, Janak PH. 2018. Ventral pallidum encodes relative reward value earlier and more robustly than nucleus accumbens. Nat Commun. doi:10.1038/s41467-018-06849-z

Pasupathy A, Miller EK. 2005. Different time courses of learning-related activity in the prefrontal cortex and striatum. Nature 433:873–876. doi:10.1038/nature03287

Redish AD. 2016. Vicarious trial and error. Nat Rev Neurosci 17:147–159. doi:10.1037/h0055482

Regier PS, Amemiya S, Redish AD. 2015. Hippocampus and subregions of the dorsal striatum respond differently to a behavioral strategy change on a spatial navigation task. J Neurophysiol 114:1399–1416. doi:10.1152/jn.00189.2015

Richard JM, Stout N, Acs D, Janak PH. 2018. Ventral pallidal encoding of reward-seeking behavior depends on the underlying associative structure. Elife 7:1–25. doi:10.7554/eLife.33107

Robbe D. 2018. To move or to sense? Incorporating somatosensory representation into striatal functions. Curr Opin Neurobiol 52:123–130. doi:10.1016/j.conb.2018.04.009

Rueda-Orozco PE, Robbe D. 2015. The striatum multiplexes contextual and kinematic information to constrain motor habits execution. Nat Neurosci 18:453–460. doi:10.1038/nn.3924

Sales-carbonell C, Taouali W, Khalki L, Pasquet MO, Petit LF, Moreau T, Rueda-orozco PE, Robbe D. 2018. No Discrete Start /Stop Signals in the Dorsal Striatum of Mice Performing a Learned Action Article No Discrete Start /Stop Signals in the Dorsal Striatum of Mice Performing a Learned Action. Curr Biol 28:3044-3055.e5. doi:10.1016/j.cub.2018.07.038

Schmitzer-Torbert NC, Redish AD. 2008. Task-dependent encoding of space and events by striatal neurons is dependent on neural subtype. Neuroscience 153:349–360. doi:10.1016/j.neuroscience.2008.01.081

Smith KS, Graybiel AM. 2016. Habit formation. Dialogues Clin Neurosci 18:33–43.

Smith KS, Graybiel AM. 2013. A dual operator view of habitual behavior reflecting cortical and striatal dynamics. Neuron 79:361–374.

Stalnaker XTA, Berg B, Aujla XN, Schoenbaum XG. 2016. Cholinergic Interneurons Use Orbitofrontal Input to Track Beliefs about Current State. J Neurosci 36:6242–6257. doi:10.1523/JNEUROSCI.0157-16.2016

Thorn CA, Atallah H, Howe M, Graybiel AM. 2010. Differential Dynamics of Activity Changes in Dorsolateral and Dorsomedial Striatal Loops during Learning. Neuron 66:781–795. doi:10.1016/j.neuron.2010.04.036

Thrailkill EA, Trask S, Vidal P, Alcalá JA, Bouton ME. 2018. Stimulus Control of Actions and Habits : A Role for Reinforcer Predictability and Attention in the Development of Habitual Behavior. J Exp Psychol Anim Learn Cogn 44:370–384.

Vandaele Y, Ahmed SH. 2020. Habit, choice, and addiction. Neuropsychopharmacol Off Publ Am Coll Neuropsychopharmacol. doi:10.1038/s41386-020-00899-y

Vandaele Y, Guillem K, Ahmed SH, Ahmed SH. 2020. Habitual Preference for the Nondrug Reward in a Drug Choice Setting. Front Behav Neurosci 14:1–9. doi:10.3389/fnbeh.2020.00078

Vandaele Y, Mahajan NR, Ottenheimer DJ, Richard JM, Mysore SP, Janak PH. 2019. Distinct recruitment of dorsomedial and dorsolateral striatum erodes with extended training. Elife 8. doi:10.7554/eLife.49536

Vandaele Y, Pribut HJ, Janak PH. 2017. Lever Insertion as a Salient Stimulus Promoting Insensitivity to Outcome Devaluation. Front Integr Neurosci 11. doi:10.3389/fnint.2017.00023

Yin HH, Knowlton BJ. 2006. The role of the basal ganglia in habit formation. Nat Rev Neurosci 7:464–476. doi:10.1038/nrn1919

Yin HH, Knowlton BJ, Balleine BW. 2006. Inactivation of dorsolateral striatum enhances sensitivity to changes in the action-outcome contingency in instrumental conditioning. Behav Brain Res 166:189–196. doi:10.1016/j.bbr.2005.07.012

Yin HH, Knowlton BJ, Balleine BW. 2004. Lesions of dorsolateral striatum preserve outcome expectancy but disrupt habit formation in instrumental learning. Eur J Neurosci 19:181–189. doi:10.1111/j.1460-9568.2004.03095.x

Yin HH, Ostlund SB, Knowlton BJ, Balleine BW. 2005. The role of the dorsomedial striatum in instrumental conditioning. Eur J Neurosci 22:513–523. doi:10.1111/j.1460-9568.2005.04218.x

