## Supplemental Figures for "Unveiling the neural correlates of habit in the dorsal striatum"

**Vandaele Y.[1], Janak P.H.[2,3]**

[1]Center for Psychiatric Neurosciences, Lausanne University Hospital – CHUV, Prilly, Switzerland. [2]Department of Psychological and Brain Sciences, Krieger School of Arts and Sciences, Johns Hopkins University, Baltimore MD 21218, USA. [3]The Solomon H. Snyder Department of Neuroscience, Johns Hopkins School of Medicine, Johns Hopkins University, Baltimore MD 21205 USA

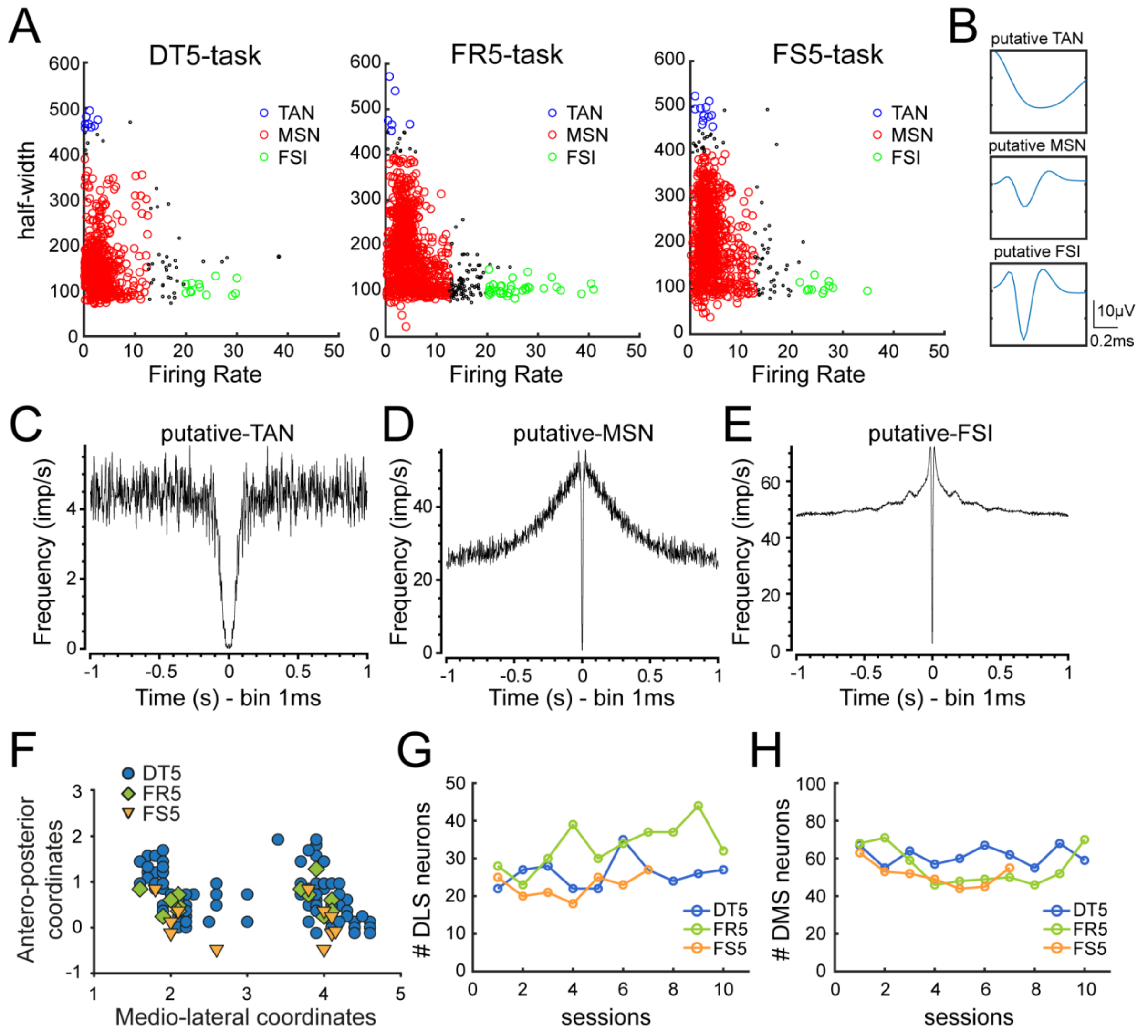


**Figure 3 – Supplement 1. Isolation of medium spiny neurons and recording summary.** (A) Scatter plots of firing rate and half-valley width in the DT5 (left), FR5 (middle) and FS5 (right) data sets. (B) Example of the average waveform of putative-TAN (top),-MSN (middle) and –FSI (bottom). (C) Autocorrelograms of representative putative-TAN (left), putative-MSN (middle) and putative-FSI (right). (D) Antero-posterior and medio-lateral coordinates of electrode placements in the DT5 (circle), FR5 (diamond) and FS5 (triangles) datasets. (G-H) Number of recorded units per session in DLS (G) and DMS (H).in the DT5, FR5 and FS5 data sets.
